# Age-related erosion of X chromosome inactivation in human tissues

**DOI:** 10.64898/2025.12.09.693282

**Authors:** Clarissa Rocca, Björn Gylemo, Mélise Edwards, Zam Cing, J. Raphael Gibbs, Colm E. Nestor, Alex R. DeCasien

## Abstract

Age-related diseases often show sex differences, yet their molecular bases remain unclear. Animal models suggest that age-related disruption of X-chromosome inactivation (XCI) occurs in female mice. We test whether this phenomenon extends to humans using bulk and single-cell datasets. We find that age-dependent escape from XCI also occurs in human females, particularly among genes at the distal ends of the X-chromosome and those involved in genome stability. These findings provide preliminary evidence that XCI erosion represents a human female-specific aging process.

## Introduction

Sex differences are a hallmark of many aging-related diseases, yet the molecular mechanisms underlying these differences remain poorly understood. Recent studies in mice have identified a female-specific age-related mechanism in which X chromosome inactivation (XCI) becomes progressively disrupted with age ^1,2^. Specifically, analyses of allele-specific expression (ASE) in *Xist*-mutant mouse models revealed that genes normally subject to XCI in young females – expressed exclusively from the active X chromosome (X_a_) – can escape silencing and become transcribed from both the active and inactive X chromosomes (X_a_ and X_i_) in aged females ^1,2^. Whether similar age-associated changes in XCI occur in humans remains largely unknown. Progress has been limited by two major challenges: i) the rarity of individuals with non-mosaic (i.e., completely skewed) XCI (nmXCI), which is required to accurately assess X-linked ASE in bulk RNA-seq data; and ii) obstacles with measuring ASE in sparse and small single cell RNA-seq datasets.

Here, we address these limitations by examining two complementary datasets. First, recent work identified female bulk RNA-seq samples in the Genotype Tissue Expression (GTEx) project with varying levels of naturally occurring XCI skew ^3–5^. These data allow for identification of samples with highly skewed nmXCI, which enable direct measurement of allelic expression (AE) – the proportion of transcripts derived from X_i_ versus X_a_ – per gene and sample ^4,5^. Age variation across these samples therefore supports our preliminary investigation into age-related XCI dynamics across human tissues. We refer to these samples collectively as the “bulk nmXCI” dataset and use them in three complementary analyses: “prior consensus”, “matched tissue”, and “linear modelling”. Second, novel tools for assessing allele-specific expression in single cell RNA-seq data have been developed and applied to new large single cell datasets, including the Asian Immune Diversity Atlas (AIDA) ^6,7^. Although ASE measurements remain sparse at the single-cell level, these data allow assessment of age-associated changes in XCI escape across substantially larger numbers of individuals because analysis is not restricted to those with extreme XCI skewing. We refer to these data as the “single-cell ASE” dataset.

Together, these complementary approaches provide the first opportunity to evaluate age-associated changes in XCI escape across human tissues and cell types. Our analyses suggest that, as observed in mice, escape from XCI increases with age in human females, particularly among genes located near the distal ends of the X chromosome and those involved in genome stability. These findings raise the possibility that progressive erosion of XCI contributes to female-biased outcomes in aging-related diseases and disorders ^1^, although changes in XCI could also reflect mechanisms of resilience ^2^.

### Many samples in the GTEx show highly skewed XCI

We collated per-gene, per-sample ASE measurements from two recent analyses of the GTEx data ^4,5^ (Methods). Allelic expression (AE) was calculated as |0.5−[reference reads/total reads]|. To facilitate interpretation, we quantified escape from XCI as 1−AE (which ranges from 0.5 to 1), such that larger values (near 1) indicate greater escape from XCI, while smaller values (near 0.5) indicate greater silencing. For each individual and tissue, we also calculated an XCI mosaicism score as the median allelic expression (1-AE) across N=263 consensus silenced genes (Methods) (Figure 1A). Because these genes are expected to be expressed exclusively from X_a_, higher XCI mosaicism scores indicate mosaic XCI, whereas lower XCI mosaicism scores indicate increasing XCI skewing ^4,5^.

**Figure 1.**
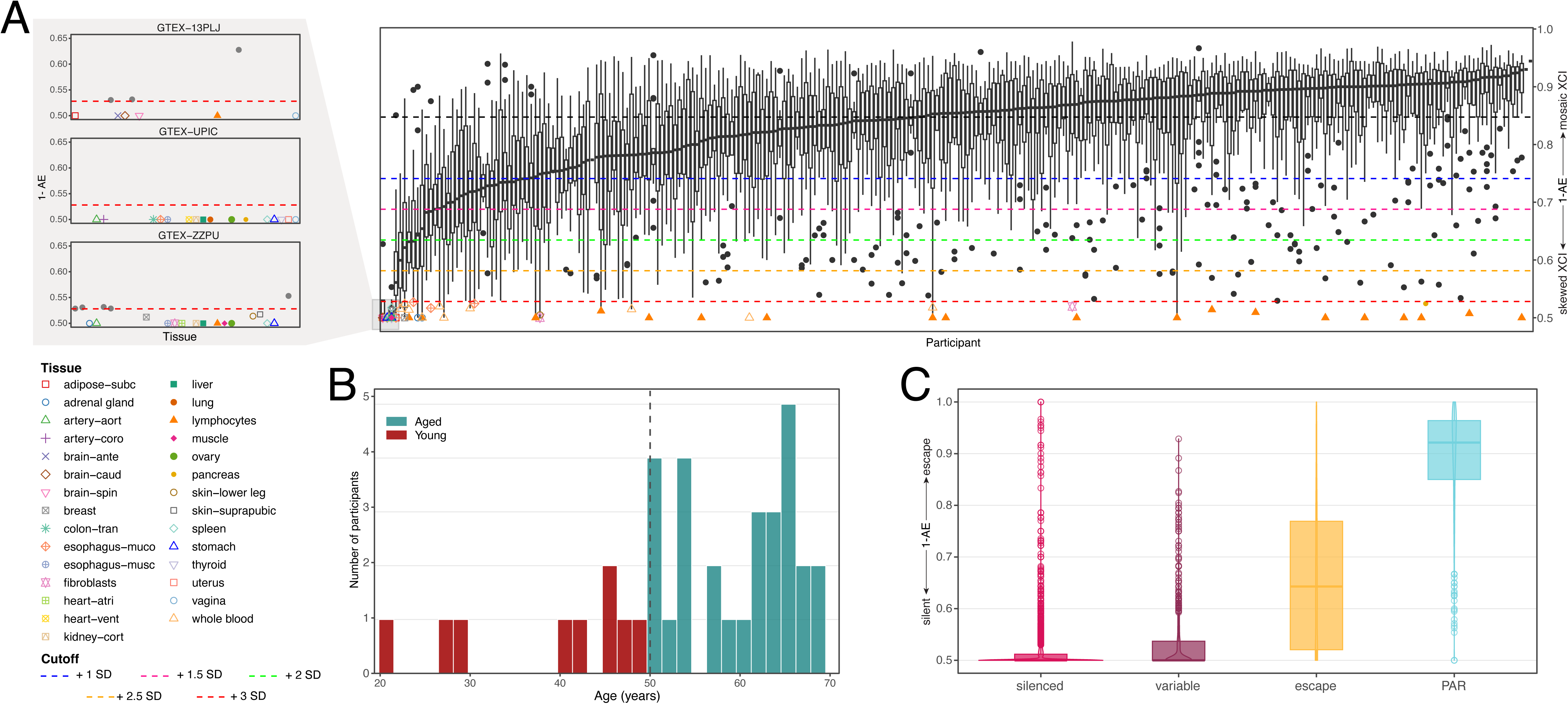
Identifying nmXCI samples in the bulk GTEx data **A.** Boxplots showing XCI mosaicism (y-axis) per individual (x-axis) and tissue (individual points). Higher values indicate mosaic (non-skewed) XCI, while lower values indicate greater XCI skewing. All points for the N=81 samples that were considered as highly skewed (<=3 SDs of the median) – and were therefore included in the bulk nmXCI dataset – are colored and shaped according to their corresponding tissue (see legend). Three individuals with highly skewed XCI across almost all tissues (identified by Gylemo and colleagues ^5^) are shown in zoomed in plot to the left. **B.** Bar plot showing age distribution of individuals included in the nmXCI dataset. The dashed line represents the 50 year cutoff for age groupings (young vs. aged). **C.** Boxplots and overlaid violin plots showing gene-level allelic expression (1-AE) (y-axis) across bulk nmXCI samples for genes with different prior XCI status classifications (x-axis). Genes without prior classification are not shown. PAR = pseudoautosomal region.

Samples exhibiting nmXCI were identified as those with XCI mosaicism scores at least three standard deviations below the population median (median=0.848, SD=0.106, cutoff=0.528) (Methods) (Figure 1A). This stringent threshold corresponds to near-complete XCI skewing and identified N=81 nmXCI female samples from N=37 unique individuals spanning 29 tissues, including three individuals previously reported to exhibit nmXCI across nearly all sampled tissues ^5^ (Figure 1A). These samples ranged in age from 21-69 years (Figure 1B) and displayed the expected allelic expression values across genes with different XCI states (Figure 1C), providing a unique opportunity to investigate age-related differences in gene-level escape from XCI in human tissues.

### Consensus silenced genes become reactivated in aged human females in the bulk nmXCI dataset

We first asked whether genes previously classified as constitutively subject to X chromosome inactivation (XCI) exhibit evidence of age-associated escape in the bulk nmXCI dataset. To do so, we leveraged consensus lists of silenced genes compiled by Balaton and colleagues ^8^ and Tukiainen and colleagues ^3^, which together identified 263 genes consistently subject to XCI across tissues and studies (Methods). For this analysis, females in the bulk nmXCI dataset were classified as “young” (<50 years) or “aged” (>=50 years), and genes were classified as escaping XCI (1−AE≥0.6) or silenced (1−AE<0.6) following the original publications ^5^ (Methods). Of the 263 consensus silenced genes, 104 (40%) had sufficient ASE data in both age groups for analysis (Methods).

Most analyzed genes (63/104, 60.6%) remained silenced (1-AE<0.6) across all samples in both age groups, indicating stable maintenance of XCI (Figure 2A). A small number of genes (4/104, 3.8%) exhibited evidence of escape in at least one sample from both age groups (Figures 2A, 2B). However, 37/104 (35.6%) genes showed age-associated differences in XCI status. The majority of these genes (30/37, 81.1%) exhibited escape from XCI (1-AE≥0.6) in at least one older sample but remained silenced in all younger samples (1-AE<0.6) (Figures 2A, 2B). Importantly, evidence of age-associated escape was observed repeatedly across independent individuals. *CHRDL1*, *HEPH*, *MPP1*, *PLXNA3*, *SSR4*, and *TCEAL4* each escaped XCI in two distinct older individuals, while *ZNF449* escaped in three older individuals (Figure 2B), suggesting that these events are unlikely to reflect sample-specific artifacts. In contrast, only 7/37 (18.9%) genes showed the opposite pattern, escaping in one young sample (1-AE≥0.6) but remaining silenced in all older samples (1-AE<0.6) (Figures 2A, 2B).

**Figure 2.**
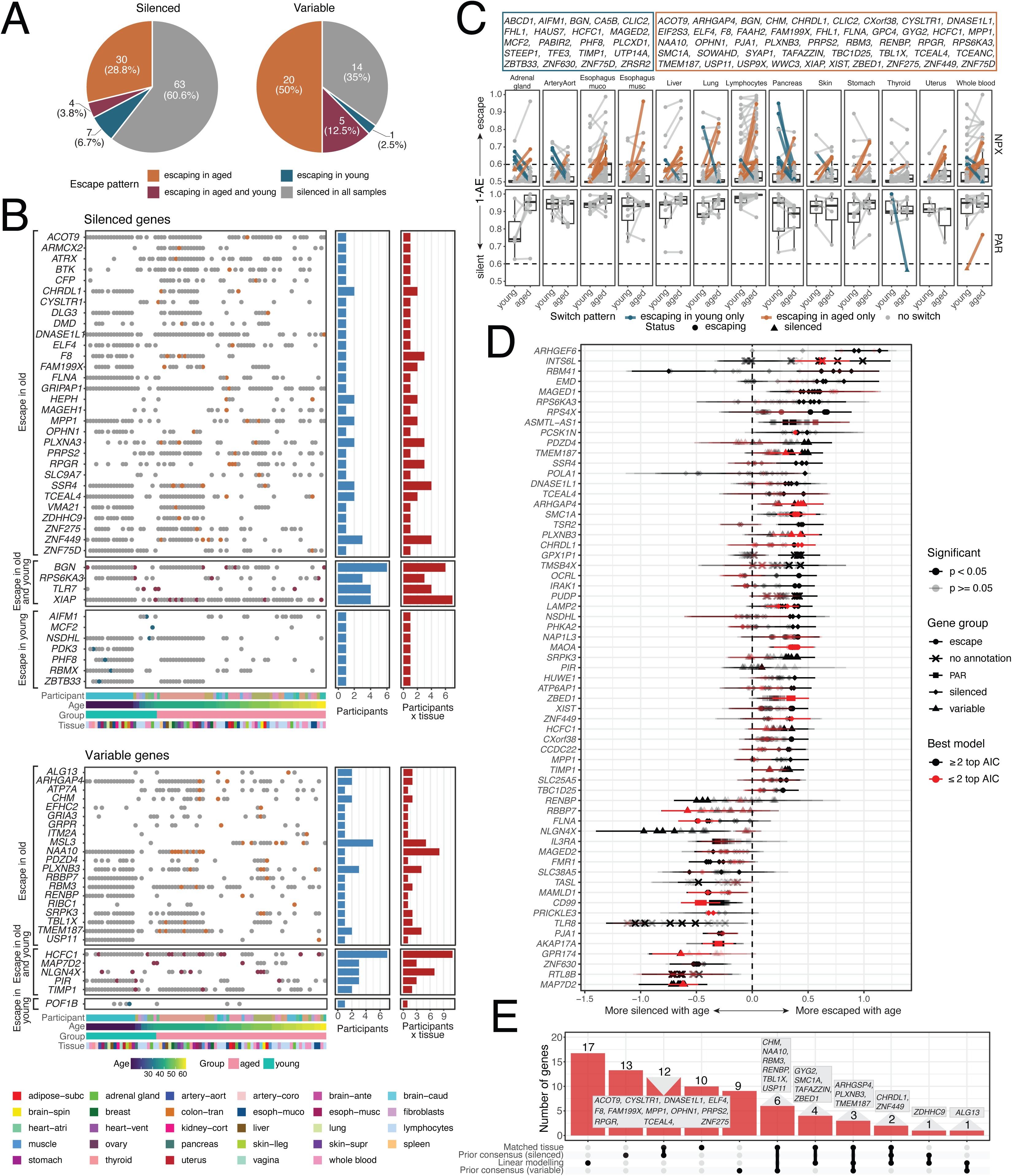
Analysis of the bulk nmXCI dataset **A.** Pie charts showing the results from the prior consensus analysis. Genes previously classified as silenced (left) or variable (right) in two published consensus lists are broken down by their age-related escape patterns in the bulk nmXCI dataset (see legend). **B.** Left: For each gene (y-axis), the XCI status (silenced if 1-AE<0.6; escaping if 1-AE>=0.6) is shown per sample (x-axis). Colors match those in Figure 2A. Metadata per sample is shown below each plot (including participant, age, age group, and tissue; see legend). Right: Bar plots showing the number of unique individuals (blue) or the total number of samples (individuals x tissues; red) showing escape from XCI per gene. Top: Genes previously classified as silenced. Bottom: Genes previously classified as variable escape. **C.** Scatterplots of allelic expression (1-AE) in young or aged individuals with each of 13 tissues. Each point represents a gene with matched data across age groups (connected by lines) (the maximum 1-AE value per age group x tissue is shown). Point and line colors and point shapes indicate age-dependent XCI status (see legend). Esophagus-muco = esophagus mucosa. Esophagus-musco = esophagus muscularis. ArteryAort = aortic artery. **D.** Dot plot showing the age effect estimate (points) and standard error of the estimate (lines) (x-axis) for each of 32 models run per gene (y-axis). Only genes where at least 1 model showed a significant age effect (p<0.05) are shown. The shape of the points represents their consensus XCI status (see legend). The color of the points represents whether the model was among the best fit models (dAIC<=2, red) or not (dAIC>2, black). Transparency of the points represents whether age was significant (p<0.05, opaque) or not (p>=0.05, transparent) in the model. **E.** Upset plot showing the numbers of genes showing age-related increases in escape from XCI in each analysis of the bulk nmXCI dataset. Genes showing mixed patterns within or across analyses are not included. Linear modelling results include best fit models only (Methods).

We next examined N=130 genes previously classified as exhibiting variable escape from XCI. Only 40/130 (30.8%) genes had sufficient data for analysis. Consistent with the results for silenced genes, evidence of escape was more common among older females. While 14/40 (35.0%) genes remained silenced across all samples, half of the analyzed genes (20/40, 50.0%) exhibited escape exclusively among older samples (Figure 2A). Again, multiple genes show age-associated escape in multiple distinct individuals (e.g., *ARHGAP4* and *TMEM187* in two, *PLXNB3* in three) (Figures 2A, 2B). By comparison, only 5/40 (12.5%) showed escape in both age groups and a single gene (1/40, 2.5%) exhibited escape exclusively among younger samples (Figure 2A). Together, these analyses provide initial evidence that aging may lead to increased escape among typically silenced genes, mirroring observations from aged mouse models, and that age-related variation may contribute to calls of “variable” XCI status.

### Tissue-matched analyses show age-dependent escape from XCI in the bulk nmXCI dataset

In the matched tissue analysis, we examined tissue-specific escape from XCI in the bulk nmXCI dataset. This analysis focused on gene x tissue combinations for which data was available in both the young and aged samples and considered all X chromosome genes regardless of their “consensus” XCI status. Across 227 unique X-linked genes and 13 tissues, we classified 897 gene x tissue combinations as either escaping XCI (1-AE≥0.6) or silenced (1-AE<0.6) based on allele-specific expression patterns in the young and aged samples (Methods). The majority of gene x tissue combinations were classified consistently across age groups: 72.8% (653/897, 185 unique genes) were silenced in both young and aged females, and 17.4% (156/897, 39 unique genes) escaped XCI in both age groups (Figure 2C). However, 9.8% (88/897; 63 unique genes) showed evidence of age-dependent changes in XCI status. Similar to the prior consensus analysis, most of these age-dependent cases (66/88, 75%; 47 unique genes) reflected a gain of XCI escape in aged females (i.e., these genes were silenced in the young samples but escaped XCI in at least one aged sample) (Figure 2C). Some genes showed recurrent XCI escape gains in aged females, including *NAA10* (4 tissues from 1 individual) and *TMEM187* (2 tissues from 2 individuals) (Figure 2C). The tissues most commonly showing gain of XCI escape with age were the esophagus mucosa (9 genes), esophagus muscularis (6 genes), liver and lung (5 genes each) (Figure 2C). Only 22/88 (25.0%; 21 unique genes) of the age-dependent gene x tissue combinations showed the opposite pattern, namely, escaping in the young samples but silencing in the aged samples (Figure 2C). Five genes showed mixed patterns (i.e., loss or gain of escape) across tissues (*BGN*, *CLIC2*, *FHL1*, *HCFC1*, *ZNF75D*) (Figure 2C), suggesting interindividual variability in escape that is not driven by aging.

### Linear modelling confirms age-dependent escape from XCI in the bulk nmXCI dataset

We next assessed the robustness of the observed age-associated increases in XCI escape using a complementary linear modeling approach. For each gene in the bulk nmXCI dataset, we modeled allelic expression (1−AE) as a function of age while accounting for combinations of three technical covariates (XCI mosaicism, uncertainty in the mosaicism estimate, and total allele count) and two biological covariates (individual and tissue) (Methods). These covariates were selected based on variance partitioning analyses, which showed that gene-level variation in allelic expression is influenced by both technical and biological factors. Across all samples, XCI mosaicism explained the largest proportion of variance (mean = 40.7%), followed by individual (6.6%), tissue (3.5%), age (1.6%), allele count (1.5%), and mosaicism uncertainty (1.3%), with the remaining 44.9% attributable to residual variation (Supplementary Figure 1).

As expected, technical effects were reduced among highly skewed nmXCI samples, where allelic expression is most informative for assessing escape from XCI. In this subset, variance explained by XCI mosaicism decreased to 12.2%, whereas the contributions of individual (18.0%), tissue (22.8%), age (5.2%), allele count (4.4%), and mosaicism uncertainty (13.0%) increased, and residual variation declined to 24.8% (Supplementary Figure 1). To ensure that our conclusions were not driven by model specification, we fit all 32 possible combinations of these five covariates for each gene. Model fit was evaluated using Akaike Information Criterion (AIC) (Methods). Technical covariates were frequently retained among the best-supported models, underscoring the importance of accounting for these factors when modeling gene-level allelic expression.

Despite accounting for these factors, evidence for age-associated increases in XCI escape remained widespread (Supplementary Table 1). Across all fitted models, 44 genes exhibited increased escape with age (effect of age [βage] > 0, nominal p < 0.05), including 11 genes supported by the best-fitting models (Figure 2D; Supplementary Table 1). More than half of these genes (25/44) also showed age-associated changes in categorical XCI status. 3/44 genes escaped XCI across all samples (1-AE≥0.6) but nevertheless exhibited increasing relative X_i_ expression with age (*RPS4X*, *PUDP*, and *ASMTL-AS1*). An additional 16/44 genes remained classified as silenced (1-AE<0.6) yet showed modest age-associated increases in relative X_i_ expression, suggesting progressive erosion of XCI that did not cross the threshold for categorical escape. Fewer genes exhibited evidence of increased silencing with age (βage < 0, nominal p < 0.05; 20 genes across all models and 8 genes in the best-fitting models) (Figure 2D; Supplementary Table 1). 6/20 of these genes showed age-associated changes in categorical XCI status. Two genes – *AKAP17A* and *IL3RA* – both located within the pseudoautosomal region (PAR), escaped XCI across all samples (1-AE≥0.6) but showed decreasing relative X_i_ expression with age (Supplementary Table 1), suggesting age-related erosion of PAR gene escape. An additional 4/20 genes remained classified as silenced (1-AE<0.6) while exhibiting modest reductions in relative X_i_ expression (Supplementary Table 1). Together, these results indicate that age-associated increases in XCI escape are substantially more common than age-associated increases in XCI silencing.

We next compared the results across the prior consensus, matched tissue, and linear modeling (best fit models only) approaches (Supplementary Table 2). Across all analyses of the bulk nmXCI dataset, we identified 130 unique genes showing age-dependent escape from XCI. Of these, 78/130 (60%) genes exclusively exhibited greater XCI escape in aged individuals, while 36/130 (27.7%) genes exclusively exhibited greater XCI escape in younger individuals (Figure 2; Supplementary Table 2). 16/130 (12.3%) genes showed more and less escape with age within the same analysis or across different analyses (Figure 2; Supplementary Table 2).

Despite their distinct assumptions, all three identified overlapping sets of genes exhibiting age-associated increases in XCI escape (Figure 2E). The greatest overlap was observed between the prior consensus and matched tissue analyses (21 genes), while 9 genes overlapped between the matched tissue and linear modeling approaches and 7 genes overlapped between the prior consensus and linear modeling approaches (Figure 2E). Notably, five genes were independently identified by all three analyses: the variably escaping genes *ARHGAP4*, *PLXNB3*, and *TMEM187*, and the consensus-silenced genes *CHRDL1* and *ZNF449* (Figure 2E). Together, these results indicate that age-associated increases in XCI escape are reproducible across multiple analytical approaches.

Interestingly, *XIST*, which is typically expressed exclusively from X_i_, showed evidence of age-associated increases in biallelic expression in both the matched tissue analysis and the linear modeling framework (although not in the best-fitting models). These findings suggest a reduction in allelic imbalance with age and are consistent with broader age-associated changes in the maintenance of XCI.

### Erosion of XCI may impact chromatin maintenance and glutamate signaling

The 78 genes that exclusively showed greater escape from XCI in the aged samples were enriched for seven pathways (g:SCS < 0.1) spanning chromatid cohesion (two terms: g:SCS = 4.85E-02, g:SCS = 4.85E-02; relevant genes = *NAA10*, *ATRX*), glutamate signaling (g:SCS = 6.48E-02; *DLG3, GRIPAP1, OPHN1*), development (hand apraxia, g:SCS = 8.12E-03; *MECP2, PLP1, SMC1A*), and location on the X chromosome (g:SCS = 1.06E-39; g:SCS = 1.07E-29; g:SCS = 2.26E-04) (Figure 3A) (Methods). These findings suggest that age-related changes in XCI may influence chromatin dynamics and genomic stability. In contrast, the 36 genes that exclusively showed escape in young individuals were enriched for five pathways (g:SCS < 0.1), including three that reflected their location on the X chromosome (g:SCS = 4.51E-21; g:SCS = 3.24E-09; g:SCS = 8.25E-02) and two related to development (aplasia, g:SCS = 4.98E-02; phalanx abnormality, g:SCS = 9.95E-02) (Figure 3A). Collectively, these results suggest that age-related relaxation of XCI affects pathways involved in epigenetic regulation, chromatin cohesion and maintenance, and receptor signaling, thereby impacting multiple functional systems.

**Figure 3.**
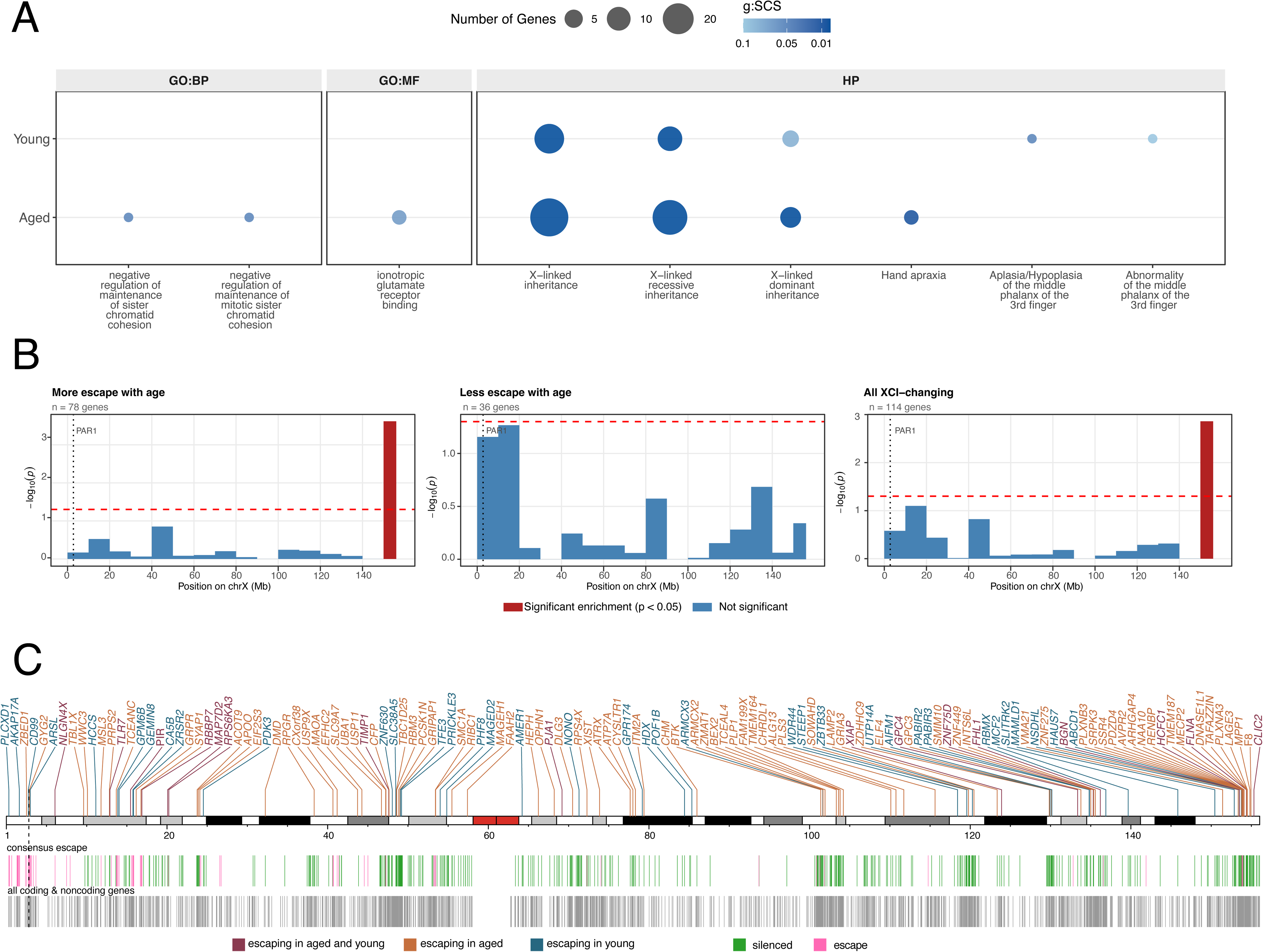
Positional and functional enrichment of genes with age-related changes in XCI status **A.** Dotplot results for functional enrichment of genes showing exclusively greater (N=78; “aged”) or lesser (N=36, “young”) escape from XCI with age. Gene ontology (GO) pathways are listed along the X axis and data sources (CORUM, biological processes [GO:BP], molecular function [GO:MF], human phenotype [HP], and microRNA target interactions [miRNA]) are highlighted above their respective GO pathways. Dot size corresponds to the number of genes and color represents the g:SCS (lower values are darker blue). **B.** Histograms show the genomic distribution of X-linked genes exhibiting exclusively increased escape with age (N=78, left), exclusively decreased escape with age (N=36, middle), and their union set (N=114, right). The X chromosome was divided into 10-Mb bins, and enrichment was assessed using one-sided Fisher’s exact tests relative to all annotated X-linked genes. Red bars indicate significant enrichment (*p* < 0.05), blue bars indicate non-significant bins, the dashed red line marks the significance threshold, and the dotted black line indicates the PAR1 boundary. **C.** Locations of genes showing age-related changes in XCI status across the human X chromosome. The X chromosome ideogram (hg38) with cytobands (from UCSC) is shown: the red area represents centromeric regions; the vertical dashed line indicates the PAR1 boundary. The union set of 130 unique genes from the analyses of the bulk nmXCI dataset and their locations are plotted above the X chromosome. The gene text color represents the pattern of age-related change observed (see legend). Directly underneath the chromosome ideogram, the colored lines indicate the location of genes assigned as either silenced or escaping XCI in either consensus list (see legend). The gray lines below denote the locations of all coding and noncoding genes along the X chromosome.

### The distal Xq end of the X chromosome appears vulnerable to XCI erosion

To understand the positional attributes of age-dependent escape genes, we performed positional enrichment analysis across the X chromosome (Methods). The 114 unique genes showing consistent age-dependent escape in the bulk nmXCI dataset (either exclusively escaping in young or in aged across analyses) were enriched at the distal Xq region (Figures 3B, 3C), with 22/114 (19.3%) of the unique age-dependent escape genes located within the last 10 Mb of the X chromosome (Fisher’s test for that 10 Mb window: OR=2.43, p=7.51E-04, all other windows p>0.05; binomial test for that 10 Mb window: p=1.12E-03; all other windows p>0.05) (Figures 3B, 3C). This finding was driven by genes that show more escape with age, of which 18/78 (23%) were located within the last 10 Mb of the X chromosome (Fisher’s test for that 10 Mb window: OR=3.04, p=2.41E-04, all other windows p>0.05; binomial test for that 10 Mb window: p=3.4E-04; all other windows p>0.05) (Figures 3B, 3C), while genes that show greater silencing with age were not enriched in any bin (Figures 3B, 3C). This is consistent with recent work showing age-dependent changes in XCI escape and chromatin accessibility at the distal ends of the X chromosome in mice ^1^, and suggests that aging is likely to disrupt compaction and silencing at the ends of the X chromosome.

### Single cell ASE data from hundreds of females demonstrate increased escape from XCI with age

In our single cell analysis, we leveraged allele-specific expression profiles from 276 female donors (19-71 years; mean = 42.2) in the Asian Immune Diversity Atlas (AIDA) ^6,7^ (Methods) to examine escape from X-chromosome inactivation at unprecedented resolution. The large sample size allowed us to treat both age and the degree of XCI escape (relative X_i_ expression) as continuous variables, enabling finer-grained modeling than was possible with bulk nmXCI dataset.

Using linear models, we examined the relationship between age and relative X_i_ expression (minor allele ratio [MAR]) for 203 genes in the single-cell ASE dataset (139 genes had sufficient data) (Methods). There were no genes that showed a negative relationship between MAR and age, while seven non-par X chromosome (NPX) genes showed a positive relationship between MAR with age (nominal p<0.05) (Figure 4). Of these, five genes (*BEX4, CYBB, HMGB3, OGT, SNX12*) were classified as silenced in the consensus lists, while two genes (*SYAP1, TCEANC*) were classified as escaping XCI (Figure 4). This suggests that age-related XCI erosion may increase expression from X_i_ and/or decrease expression from X_a_ regardless of the XCI status of a given gene. Notably, both escape genes *SYAP1* and *TCEANC* also showed age-related escape from XCI in the matched tissue analysis of the bulk nmXCI dataset from GTEx (Figure 2C), providing independent support across datasets and analytical approaches. Although exploratory, these single-cell analyses provide independent support for age-associated increases in X_i_ expression and suggest that age-related changes in XCI may extend to both silenced and escaping genes.

**Figure 4.**
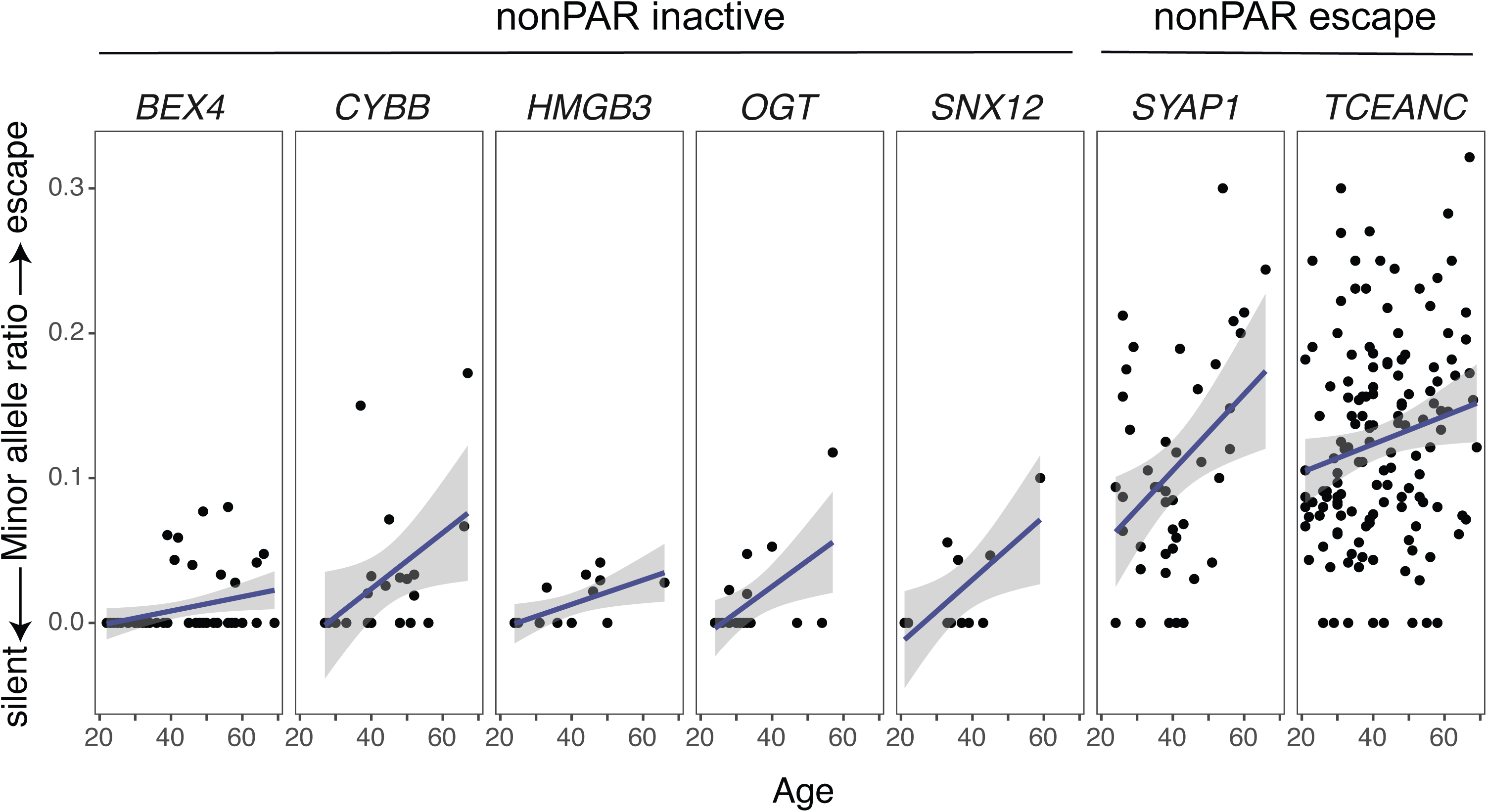
Analysis of the single cell ASE dataset Scatterplot showing the minor allele ratio (MAR, y-axis) as a function of age (x-axis) for each of the 7 genes that showed significant (p<0.05) relationships between MAR and age. Individual points represent samples, regression lines are shown in purple, and 95% confidence intervals are in grey. The XCI status of each gene is shown above the corresponding plot.

## Discussion

This study provides preliminary evidence that escape from XCI increases with age in human females, extending recent findings in mice ^1,2^. Across three complementary analyses of rare individuals with naturally occurring non-mosaic XCI, we consistently observed a bias toward increased relative expression from X_i_ in older individuals. In the prior consensus analysis, genes previously classified as constitutively silenced were more likely to exhibit escape in aged females than in younger females. The matched tissue analysis identified age-associated gains of escape across multiple tissues and genes regardless of their previously assigned XCI status. Finally, quantitative linear modeling confirmed that age-associated increases in relative X_i_ expression were more common than age-associated decreases, even after accounting for technical and biological sources of variation. Importantly, five genes (*CHRDL1*, *ZNF449*, *ARHGAP4*, *PLXNB3*, and *TMEM187*) were identified across all three approaches, providing particularly strong support for age-associated erosion of XCI at these loci.

The single-cell ASE analysis provided an orthogonal line of evidence supporting this conclusion. Although no genes survived multiple testing correction, all nominally significant associations were in the direction of increased relative X_i_ expression with age, and none showed the opposite pattern. Two genes identified in the single-cell dataset (*SYAP1* and *TCEANC*) also exhibited age-associated escape some analyses of the bulk nmXCI dataset. While these results should be interpreted cautiously given the sparsity of single-cell ASE measurements, the directional consistency across hundreds of individuals and multiple datasets provides additional support for a model in which Xi transcription gradually increases with age.

Our findings are broadly consistent with recent studies in mice demonstrating age-associated disruption of XCI ^1,2^. Several genes identified in the current study also exhibit age-related escape in mice (*CFP*, *PDK3, USP11* ^1^; *GRIPAP1*, *PLXNB3*, *SRPK3*, *PDZD4*, *NAA10* ^2^), suggesting some degree of cross-species conservation. Notably, some of these shared genes are positioned near the ends of the X chromosome in one species but not the other – for example, *GRIPAP1* lies close to the end of the mouse X chromosome (X:7,656,234-7,686,802) but is located farther from the telomere in humans (X:48,973,723-49,002,264). This pattern implies that age-related escape at these loci may relate more to their gene functions than to their chromosomal positioning alone.

One of the most striking patterns to emerge from our analyses was the positional enrichment of age-associated escape genes at the distal end of Xq. Almost twenty percent of all genes exhibiting age-dependent changes in escape were located within the final 10 Mb of the X chromosome, representing a significant enrichment relative to expectation. This finding closely parallels observations in aged mice, where chromatin accessibility and escape from XCI increase preferentially near the ends of the X chromosome ^1^. Together, these results suggest that telomeric or subtelomeric regions of the X chromosome may be particularly vulnerable to age-associated deterioration of chromatin compaction and epigenetic silencing mechanisms.

Genes exhibiting age-associated increases in escape were enriched for pathways related to sister chromatid cohesion, receptor organization, and glutamatergic signaling. These enrichments should be viewed as exploratory, but they are notable because several implicated genes participate in processes linked to genome stability and cellular aging. In particular, genes involved in chromatid cohesion, including *ATRX* and *NAA10*, may point toward a mechanistic connection between age-associated dysregulation and erosion of XCI. Age-related loss of cohesion in somatic tissues can lead to chromosomal aneuploidies and inhibit DNA repair during cell division ^9,10^, both of which are observed in aging. Whether increased escape from XCI contributes directly to aging phenotypes or instead reflects broader age-related changes in chromatin architecture remains an important question for future investigation.

Several limitations should be considered when interpreting these findings. The bulk nmXCI analyses rely on 81 highly skewed samples – 40% of which are from lymphocytes or whole blood – from 37 individuals, reflecting the rarity of naturally occurring nmXCI in human tissues. In addition, tissue availability differed across individuals, limiting power to detect tissue-specific patterns of escape. Although the convergence of three analytical approaches strengthens confidence in the results, larger cohorts of individuals with skewed XCI will be needed to establish the prevalence and magnitude of age-associated XCI erosion. Importantly, all analyses presented here are cross-sectional. As a result, we cannot directly determine whether the observed differences reflect within-individual increases in XCI escape over time or cohort-specific differences between younger and older individuals. Longitudinal studies that repeatedly measure allele-specific expression within the same individuals will therefore be essential to definitively establish the temporal dynamics of XCI erosion during human aging. Similarly, while the single-cell analysis enabled examination of many more individuals, sparse ASE measurements limited statistical power and prevented definitive identification of age-associated escape genes after multiple-testing correction.

Despite these limitations, our findings suggest that X chromosome inactivation is not a static process across the human lifespan. Instead, multiple lines of evidence indicate that expression from the inactive X chromosome increases with age, particularly among genes located near the distal end of Xq. Together with recent findings in mice, these results support a model in which age-associated erosion of XCI represents a conserved female-specific feature of aging that may contribute to interindividual variation and sex differences in age-related biology.

## Methods

### Datasets

#### Consensus lists of XCI status per gene

Balaton and colleagues ^8^ created a catalog of X chromosome genes and their consensus XCI status by integrating results from three studies of human-mouse hybrid cell lines, X-linked SNPs in microarray data, or CpG island methylation data. Genes were placed into 1 of 8 categories depending on concordance across studies, resulting in consensus XCI status for 639 genes. These methods are described further in Balaton et al. ^8^.

Tukiainen and colleagues ^3^ also provided a catalog of X chromosome genes and their consensus XCI status. They similarly integrate results from two studies of human-mouse hybrid cell lines and X-linked SNPs in microarray data, but also integrate these with their original analyses of: i) female-biased expression across human tissues in the GTEx; ii) allele-specific expression from cross-tissue bulk RNAseq of one individual with nmXCI in the GTEx; and iii) allele-specific expression from a small single cell RNAseq dataset of four females ^3^. They suggest new evidence for escape for seven genes (*ATP6AP2*, *RP13-216E22.4*, *CHM*, *RBMX2*, *FHL1*, *MAP7D3*, *CLIC2*), and we encoded them as such.

We identified N=263 consensus silenced genes across datasets (Balaton “S” AND Tukiainen “Inactive”), N=504 union silenced genes (Balaton “S” OR Balaton “Mostly S” OR Tukiainen “Inactive”), N=22 consensus escape genes (Balaton “E” AND Tukiainen “Escape”), N=94 union escaping genes (Balaton “E” OR Balaton “Mostly E” OR Tukiainen “Escape”), N=26 consensus variable genes (Balaton “VE” AND Tukiainen “Variable”), and N=130 union variable genes (Balaton “VE” OR Balaton “Mostly VE” OR Balaton “Discordant” OR Tukiainen “Variable”).

#### Bulk allele specific expression (bulk RNAseq from GTEx)

Tissue-specific XCI status for X chromosome genes were collected from two recent analyses of female GTEx donors. One study identified and characterized XCI patterns across 30 tissues in three females with consistent non-mosaic (i.e., highly skewed) X chromosome inactivation ^5^. Another study characterized XCI patterns across 35 tissues in 264 females ^4^. The former analyses provided more gene-level information since it used both whole genome and exome sequencing to call variants, while the latter used whole exome sequencing only ^4,5^. Both studies quantified allelic expression (AE) for each gene and sample as the absolute deviation from balanced expression at heterozygous sites (AE = |0.5 − [reference reads / total reads]|), where AE = 0 indicates biallelic expression and AE = 0.5 indicates complete monoallelic expression ^4,5^.

To identify female samples with “highly skewed XCI”, we followed previous work ^4,5^ and calculated an XCI mosaicism score as the median allelic expression (1-AE) across N=263 consensus silenced X chromosome genes for each individual x tissue sample. We identified N=81 highly skewed samples (representing N=37 unique individuals and 29 tissues) with XCI mosaicism values ≤3 standard deviations of the median XCI mosaicism across all samples (median=0.848, SD=0.106, cutoff=0.528), indicative of near-complete inactivation of one parental X chromosome. As expected, lymphocytes (N=20) and whole blood (N=12) were the most frequently represented sample types among highly skewed samples – consistent with the selective effects of clonal hematopoiesis ^5^ – while high skew among solid tissues was less frequent (adipose-subc=2, adrenal gland=4, artery-aort=2, artery-coro=1, brain-ante=1, brain-caud=1, brain-spin=1, breast=2, colon-tran=1, esophagus-muco=5, esophagus-musc=2, fibroblasts=2, heart-atri=1, heart-vent=1; kidney-cort=1, liver=2, lung=2, muscle=2, ovary=1, pancreas=3, skin-lleg=2, skin-supr=2, spleen=1, stomach=2, thyroid=2, uterus=2, vagina=1).

#### Single-cell allele specific expression (single-cell RNAseq from AIDA)

Tomofuji and colleagues ^6^ recently developed the *scLinaX* software to directly quantify relative gene expression from the active and inactivated X chromosomes (X_a_ and X_i_) using single-cell RNA-seq data while accounting for the sparsity and transcriptional bursting that complicate allele-specific analyses at single-cell resolution. Briefly, scLinaX first constructs pseudobulk allele-specific expression (ASE) profiles using reads overlapping informative heterozygous single nucleotide polymorphisms (SNPs). Alleles residing on the same X chromosome are then grouped using correlation analyses of these pseudobulk ASE profiles, allowing assignment of X_a_ and X_i_ identity. Individual cells are subsequently classified according to their inferred XCI state, and allele-specific information is aggregated across cells to estimate gene-level relative expression from X_i_ for each individual.

Importantly, scLinaX does not infer XCI escape from monoallelic or biallelic expression calls in individual cells. Instead, allele-specific reads are aggregated across many cells following XCI-state assignment, generating individual-level estimates of relative X_i_ expression that are substantially less sensitive to transcriptional bursting and allelic dropout than cell-level measurements. We therefore interpret the resulting values as estimates of average X_i_ contribution to gene expression within each individual rather than direct measurements of escape in individual cells. Tomofuji and colleagues applied scLinaX to the Asian Immune Diversity Atlas (AIDA) ^7^, generating estimates of relative X_i_ expression for X-linked genes across hundreds of donors.

### Analyses

#### Analyses of bulk nmXCI dataset (GTEx)

##### Prior Consensus Analysis

We combined the consensus silenced genes (N=263) obtained from Balaton and colleagues ^8^ and Tukiainen and colleagues ^3^ (Methods) with the bulk allele-specific expression (ASE) data from all highly skewed samples (<=3 SDs of median XCI mosaicism; N=81 samples; N=37 unique individuals) (Methods) to assess whether consensus silent genes show age-dependent patterns of escape (“aged”: >50 years; N=57 samples; N=28 individuals; young: <=50 years; N=24 samples; N=9 individuals). We found N=104 genes that were present (i.e., had available data) in at least one young and old sample in the bulk nmXCI dataset. We then tested whether each consensus silent gene was: i) silent in all samples across both age groups (aged, young); ii) inactive in the young group but escaping (1-AE≥0.6) in at least one aged sample; iii) inactive in the aged group but escaping (1-AE≥0.6) in at least one young sample; or iv) escaping in at least one young and one aged sample. We also repeated this analysis for N=130 genes showing variable escape across both consensus gene lists (Methods). Of these, N=40 genes were also present in at least one young and old sample in the bulk nmXCI dataset.

##### Matched Tissue Analysis

We utilized the bulk nmXCI dataset to identify tissue-specific patterns of escape with age. We required gene x tissue combinations to have data from at least one individual in each age group. Some tissues (including e.g., the brain) did not have data available for at least one individual from each age group ^5^ and were therefore not analyzed.

##### Variance Partitioning

For each gene, we estimated the proportion of variance in allelic expression (1-AE) attributable to biological factors – including participant and tissue – and technical factors – including XCI mosaicism score, uncertainty in the XCI mosaicism score (interquartile range of allelic expression (1-AE) across N=263 consensus silenced X chromosome genes), and total read count. Linear mixed-effects models were fit separately for each gene, with participant and tissue modeled as random effects using the *lmer* function in the R package *lme4*. Genes with fewer than five observations, fewer than three participants, or fewer than two tissues were excluded. For fixed effects, variance explained by each predictor was estimated as the variance of the predictor multiplied by its fitted coefficient. Random-effect variance components for participant, tissue, and residual variance were extracted from the fitted model. Variance fractions were then calculated by dividing each component by the sum of all fixed-effect, random-effect, and residual variance components.

##### Linear Modelling

To identify genes showing age-dependent changes in allelic expression among nmXCI samples (N=81 samples with XCI mosaicism <= 3 SDs below the median), we fit linear models using either the *lm* function from the R package *stats* or the *lmer* function from the R package *lme4*. The outcome variable in all models was allelic expression (1-AE), and the primary predictor of interest was age, modeled as a continuous variable (years). To ensure our findings were not driven by any single modeling decision, we fit 32 models per gene representing all possible combinations of five covariates: participant, tissue, XCI mosaicism, uncertainty in the XCI mosaicism score, and total read count. For genes where participants had data across multiple tissues, we fit a linear mixed model with a random intercept for individual (1 | participant); otherwise, we used a standard linear model. All continuous predictors used in the models were mean centered and scaled. To identify the best-fitting model per gene, we used the Akaike Information Criterion (AIC). Models with ΔAIC ≤ 2 of the best fit model (with the lowest AIC value) were considered equivalent.

#### Functional enrichment

We tested for functional enrichment of genes showing age associated changes in XCI in the bulk nmXCI dataset (for N=78 unique genes escaping in aged individuals; and for N=36 unique genes showing escape in young individuals). Functional enrichment analyses were run using the *gost* function in the R package *gProfiler2* (background set = “only annotated genes”) ^11^. Pathways with g:SCS<0.1 were considered significant. The g:SCS is the default method for multiple test correction in g:Profiler.

#### Positional enrichment

We tested for positional enrichment along the X chromosome of genes showing age associated changes in XCI in the bulk nmXCI dataset (N=78 unique genes escaping in aged individuals; N=36 unique genes showing escape in young individuals; and N=114 union genes). Locations for target genes – as well as for all coding and noncoding genes on the X chromosome – were obtained using the *getBM* function in the R package *biomaRt* ^12^. We visualized the positions of genes using the *karyoploteR* R package ^13^.

To determine whether age-dependent escape genes from analyses of the nmXCI dataset show preferential localization along the X chromosome, we tested for positional enrichment using a bin-based framework whereby the X chromosome was divided into 16 equal genomic bins (10 Mb each). We tested for positional enrichment of age-dependent escape genes along the X chromosome within each 10 Mb window using: i) binomial tests (binom.test function in R package *stats*) of the number of age-dependent escape genes in the window compared to the total number of age-dependent escape genes; and ii) one-sided Fisher’s exact test (*fisher.test* function in the R package *stats*) comparing the number of age-dependent escape genes observed in the window to the number expected by chance, given the distribution of all X-linked genes across the chromosome.

#### Analyses of the single cell dataset (AIDA)

Because only gene-level summary statistics are publicly available, we obtained individual x gene-level minor allele ratio (MAR) estimates directly from Tomofuji and colleagues ^6^. Following the original study, MAR was used as a proxy for relative X_i_ expression, with larger values indicating greater transcription from X_i_. We analyzed data from 276 female donors spanning 19-71 years of age (mean = 42.2 years). For each gene, we modeled MAR as a function of age using linear regression (*lm* function in the R package *stats*) and identified genes showing age-associated changes in MAR. Models were only run if there were at least three individuals with more than one distinct age available. We obtained results for 139/203 genes included in the original dataset. Because the single-cell ASE data represent aggregated estimates of X_i_ expression rather than categorical escape calls, these analyses were used to assess age-associated trends in X_i_ transcription and to provide an independent line of evidence complementary to the bulk nmXCI analyses.

## Declarations

### Data availability

The prior consensus XCI status lists ^3,8^ and one bulk allele-specific expression (ASE) dataset analyzed in this manuscript are publicly available ^5^. The other bulk ASE dataset is available upon request from the authors ^4^. Individual-level ASE data from the AIDA single cell dataset was provided by Tomofuji and colleagues ^6^.

### Code availability

All code used in this manuscript can be found at https://github.com/coevolving-unit/aging-gtex-xci.

### Author contributions

A.R.D. designed the study. B.G. and C.E. estimated and contributed the allele-specific bulk RNAseq data. A.R.D., C.R., M.E., and J.R.G. analyzed the data. A.R.D. and C.R. wrote the first draft manuscript. All authors reviewed and revised the manuscript.

## Acknowledgements

We thank Dr. Yoshihiko Tomofuji for generously sharing their analyses of the AIDA dataset.

This work is supported by the Intramural Research Program of the National Institute on Aging (NIA). This research was supported [in part] by the Intramural Research Program of the National Institutes of Health (NIH). The contributions of the NIH author(s) are considered Works of the United States Government. The findings and conclusions presented in this paper are those of the author(s) and do not necessarily reflect the views of the NIH or the U.S. Department of Health and Human Services.

## Ethics declaration

Competing interests: The authors declare no competing financial or non-financial interests

## Supplementary Figure Legends

**Supplementary Figure 1.**
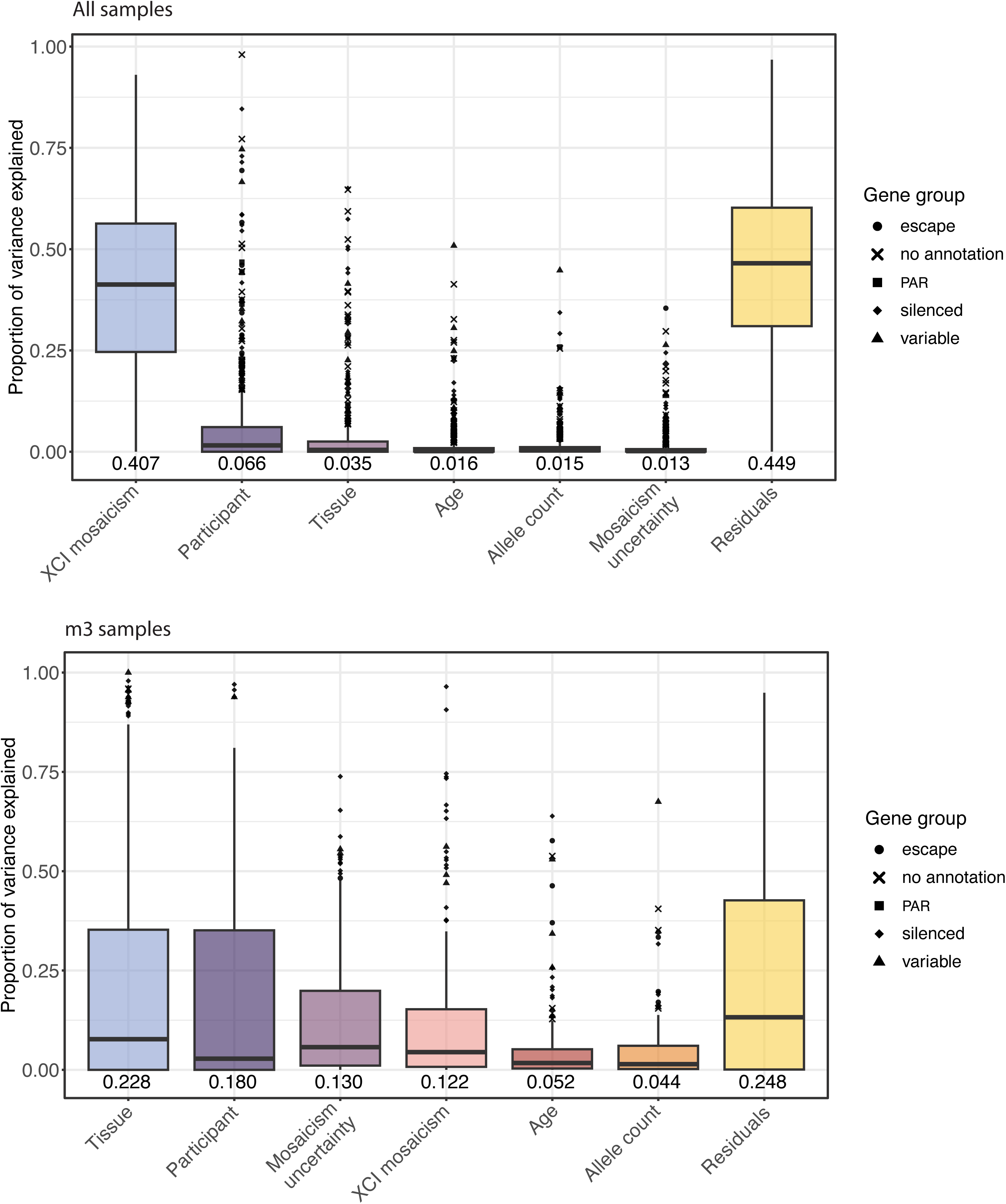
Variance partitioning of allelic expression **Top.** Boxplots of the proportion of variance in allelic expression explained per gene (y-axis) by each predictor variable (x-axis). Each gene has one point in each boxplot, and these values add up to 1. The mean proportion of variance explained by each predictor across all genes is shown below each boxplot. All samples are included. **Bottom.** Same as the top plot, but for the N=81 nmXCI samples only.

## Supplementary Table Legends

Supplementary Table 1 | Linear modelling results

For each gene, the linear modelling results across 32 models are shown. Predictors included in each model are indicated in Columns D-H. Numbers of samples, participants, and tissues are provided in Columns J-L. Model run information is in Column N (lm = linear model; lmer = linear mixed model; lmer_singular = linear mixed model with singular fit; skipped_n_ind<=3 = model was skipped because of too few individuals; failed = model failed to run). Model results (for the age effect) are in Columns P-W. Model comparisons are in Columns Y-AB).

Supplementary Table 2 | Comparison across analysis of the bulk nmXCI dataset

For each gene, the results from the prior consensus, matched tissue, and linear modelling analyses are summarized. Linear modelling results reflect those from best fit models (Methods). “Mixed” in prior consensus = escaping in aged and young samples. “Mixed” in matched tissue = mixed pattern across tissues. “No switch” = no age difference in XCI status across tissues. NA = gene was not included in the analysis.

## References

1. Hoelzl, S., Hasenbein, T. P., Engelhardt, S. & Andergassen, D. Aging promotes reactivation of the Barr body at distal chromosome regions. Nat. Aging 5, 984–996 (2025).

2. Gadek, M. et al. Aging activates escape of the silent X chromosome in the female mouse hippocampus. Sci. Adv. 11, eads8169 (2025).

3. GTEx Consortium et al. Landscape of X chromosome inactivation across human tissues. Nature 550, 244–248 (2017).

4. Goldmann, D. et al. TRiXi: A multi-target tandem repeat-based method for accurate detection of X-inactivation skewing in humans. medRxiv (2025) doi:10.1101/2025.11.24.25340860.

5. Gylemo, B., Bensberg, M. & Nestor, C. E. A whole-organism landscape of X-inactivation in humans. (2025) doi:10.7554/elife.102701.1.

6. Tomofuji, Y. et al. Quantification of escape from X chromosome inactivation with single-cell omics data reveals heterogeneity across cell types and tissues. Cell Genom. 4, 100625 (2024).

7. Kock, K. H. et al. Asian diversity in human immune cells. Cell 188, 2288–2306.e24 (2025).

8. Balaton, B. P., Cotton, A. M. & Brown, C. J. Derivation of consensus inactivation status for X-linked genes from genome-wide studies. Biol. Sex Differ. 6, 35 (2015).

9. Piazza, A. et al. Cohesin regulates homology search during recombinational DNA repair. Nat. Cell Biol. 23, 1176–1186 (2021).

10. Brooker, A. S. & Berkowitz, K. M. The roles of cohesins in mitosis, meiosis, and human health and disease. Methods Mol. Biol. 1170, 229–266 (2014).

11. Kolberg, L., Raudvere, U., Kuzmin, I., Vilo, J. & Peterson, H. gprofiler2 -- an R package for gene list functional enrichment analysis and namespace conversion toolset g:Profiler. F1000Res. 9, ELIXIR–709 (2020).

12. Durinck, S. et al. BioMart and Bioconductor: a powerful link between biological databases and microarray data analysis. Bioinformatics 21, 3439–3440 (2005).

13. Gel, B. & Serra, E. karyoploteR: an R/Bioconductor package to plot customizable genomes displaying arbitrary data. Bioinformatics 33, 3088–3090 (2017).

